# Spatiotemporal dynamics of river viruses, bacteria and microeukaryotes

**DOI:** 10.1101/259861

**Authors:** Thea Van Rossum, Miguel I. Uyaguari-Diaz, Marli Vlok, Michael A. Peabody, Alvin Tian, Kirby I. Cronin, Michael Chan, Matthew A. Croxen, William W.L. Hsiao, Judith Isaac-Renton, Patrick K.C. Tang, Natalie A. Prystajecky, Curtis A. Suttle, Fiona S.L. Brinkman

## Abstract

Freshwater is an essential resource of increasing value, as clean water sources diminish. Microorganisms in rivers, a major source of renewable freshwater, are significant due to their role in drinking water safety, signalling environmental contamination^1^, and driving global nutrient cycles^2,3^. However, a foundational understanding of microbial communities in rivers is lacking^4^, especially temporally and for viruses^5‒7^. No studies to date have examined the composition of the free-floating river virome over time, and explanations of the underlying causes of spatial and temporal changes in riverine microbial composition, especially for viruses, remain unexplored. Here, we report relationships among riverine microbial communities and their environment across time, space, and superkingdoms (viruses, bacteria, and microeukaryotes), using metagenomics and marker-based microbiome analysis methods. We found that many superkingdom pairs were synchronous and had consistent shifts with sudden environmental change. However, synchrony strength, and relationships with environmental conditions, varied across space and superkingdoms. Variable relationships were observed with seasonal indicators and chemical conditions previously found to be predictive of bacterial community composition^4,8‒10^, emphasizing the complexity of riverine ecosystems and raising questions around the generalisability of single-site and bacteria-only studies. In this first study of riverine viromes over time, DNA viral communities were stably distinct between sites, suggesting the similarity in riverine bacteria across significant geographic distances^10‒12^ does not extend to viruses, and synchrony was surprisingly observed between DNA and RNA viromes. This work provides foundational data for riverine microbial dynamics in the context of environmental and chemical conditions and illustrates how a bacteria-only or single-site approach would lead to an incorrect description of microbial dynamics. We show how more holistic microbial community analysis, including viruses, is necessary to gain a more accurate and deeper understanding of microbial community dynamics.

## Main

Bacterial diversity and composition in rivers is shaped by water temperature, day length, pH^10^, nutrients^8,9^, water residency time^10,13^, and storm events (reviewed in ^4^). Balancing these shaping forces, dispersal appears to play a large role both within^9^ and among^10‒12^ rivers, such that bacterial community similarity does not necessarily decrease with increasing geographic distance. Less is known about planktonic (free-floating) microeukaryotes in rivers, however, they appear to vary seasonally with light changes^14‒16^, with some evidence indicating the importance of algae as an energy source^15^.

In contrast to this basic characterisation of bacterial and microeukaryote community variability, little is known about the community dynamics of free-floating viruses (viroplankton) in rivers^5‒7^. River planktonic viral metagenomes (viromes) have been reported in two studies^17,18^, however, these studies had limited sample sizes and did not sample over time. Viral communities in lakes and oceans are better studied, however, these viromes are likely distinct from those in rivers given their differing hydrology and bacterial community compositions^7,10,19^. To date, there have been no large-scale studies of viroplankton composition in flowing (lotic) freshwater. As such, little is known about their community composition^5-7^ and basic questions, such as their variability throughout a year and the relative importance of dispersal and shaping forces in their community composition have gone unanswered.

Fundamental knowledge of the spatiotemporal variability of river plankton can support downstream development of improved water quality indicators. To this end, we profiled viral, bacterial, and microeukaryotic communities in rivers across differing land uses and environmental conditions. We sampled microorganisms monthly for one year from six sites in three watersheds in southwestern British Columbia, Canada (Figure 1a). For each sample, we performed metagenomic and/or phylogenetic marker gene sequencing (16S, 18S, g23 viral capsid) for DNA viruses, RNA viruses, bacteria^20^, and microeukaryotes^21^. Environmental, chemical, and biological measures were also collected^20,21^. Positive and negative controls were included, and qPCR validation of select microbial groups was performed (data not shown). Due to the lack of reference genomes available for freshwater viruses and the high complexity of the communities, we estimated dissimilarity measures among metagenomes using a reference- and assembly-free k-mer approach (Mash^22^). To diminish any effects from potential bacterial or eukaryotic contamination in the viral data, DNA and RNA viromes are represented by two datasets. The “total” dataset includes all sequence reads. The “conservative” dataset is a subset of reads selected based on similarity to known viruses (see Methods for details). Spatiotemporal comparisons were performed within and between “superkingdoms”, including viruses (DNA and RNA), bacteria, and microeukaryotes, and “environmental conditions”, including catchment area weather, river water chemical concentrations, and river water physical conditions.

**Figure 1.**
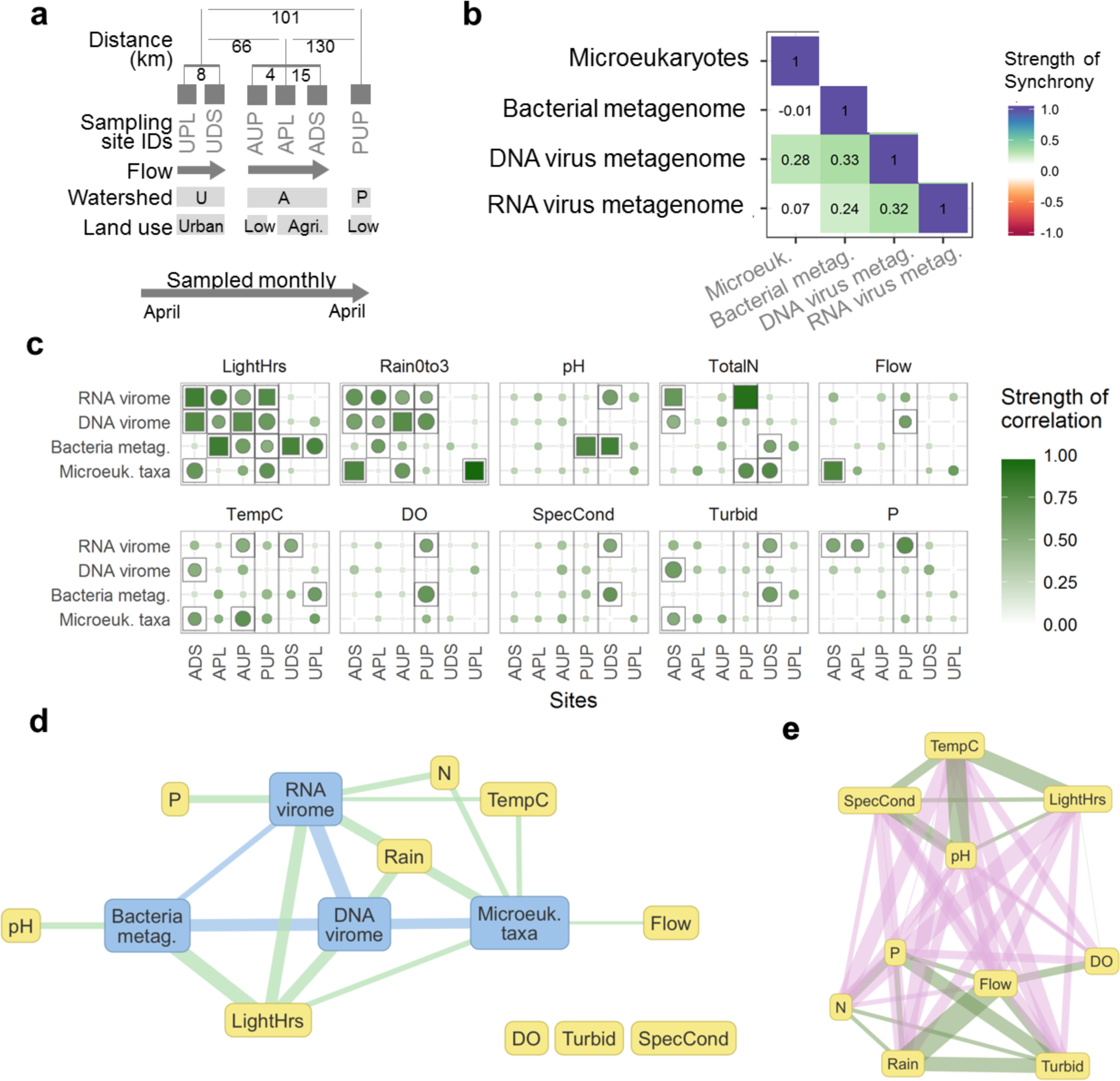
Temporal variation in viruses, bacteria, and microeukaryotes. **a**, Study design schematic of sampling sites with distances between sites, site orientation, watershed, and catchment land use. Distances are dendritic within watersheds and Euclidean between watersheds. Sites are in up- to down-stream order within watersheds. **b**, Pairwise partial Mantel tests for synchrony between viruses, bacteria and microeukaryotes, controlling for distance between sampling sites, N = 51 to 85, q < 0.0004. **c**, Correlations between microbial communities and environmental conditions per sampling site. Results are organised by environmental parameter into subplots where each row is a biological group and each column is a sampling site. Colour intensity reflects correlation strength. Filled shapes indicate the statistical significance of the correlation with squares as significant (q < 0.1) and circles not statistically significant. Size of shape corresponds to the inverse of the statistical significance (q value). Grey square outlines indicate a relationship was statistically significant without multiple test correction (p < 0.05). Grey vertical lines separate watersheds. **d**, Network of summarised correlations among microbial communities and with environmental conditions, calculated per sampling site. Nodes are environmental conditions (yellow) and microbial communities (blue). Conservative viromes were used (see methods). Edges are coloured by the nodes types they connect. Each edge represents cumulative relationships within sampling sites, both those that are statistically significant (q < 0.1) and that are strong but with lower statistical confidence (R^2^ > 0.34, p < 0.05). Edge width reflects the sum of the strengths (R^2^) of the represented correlations. Edges are only drawn if at least one statistically significant or two lower-confidence correlations were observed, to reduce artefacts from arbitrary statistical cut-off values. e, Network of correlations among environmental conditions, with edges calculated as in (d), with green edges for positive correlations and pink for negative. Nodes were arranged manually for legibility.

Across superkingdoms, hours of daylight and rainfall intensity were the most commonly correlated with community composition (Figure 1 c, d). This pattern was particularly strong where rainfall and hours of daylight were correlated (Figure 1c sites AUP, APL, ADS; Extended Data Fig. 2 b, c, d), but weak in sites where they were not (Figure 1c sites PUP, UPL, UDS; Extended Data Fig. 2 a, e, f). This is surprising as rainfall was hypothesized to have a particularly large and consistent impact on microbial communities since its intensity can affect microbial transport (both overland and within stream transport). Instead, when not confounded with overall seasonal changes (hours of daylight), rainfall was rarely significantly correlated with microbial community composition. Overall, no correlations between environmental conditions and superkingdoms were seen in all sites (Figure 1c), emphasizing the variability of river microbial community relationships with their environment.

Environmental conditions that have been reported to drive bacterial community composition were heterogeneously correlated across sites and did not extend to other superkingdoms. For example, nitrogen and phosphorous concentrations were most often correlated with RNA viruses and/or microeukaryotes but not with bacteria, and pH was only correlated with bacterial composition in two sites, despite a previous single-time-point study finding it to be a major driver^10^. Very few correlations were observed with dissolved oxygen concentration, flow intensity, specific conductivity, or turbidity. The range of correlations with environmental conditions observed across sites and superkingdoms emphasizes both the complexity and heterogeneity of riverine microbial ecosystems.

Despite inconsistent relationships with environmental conditions, viral and bacterial community compositions shifted in similar patterns over time (were “synchronous”), with the strength of synchrony varying among sampling sites (Figure 2b, Extended Data Fig. 2). Microeukaryotes had fewer synchronous relationships but were correlated with bacteria and/or DNA viruses in some sites. The lack of synchrony between microeukaryotes and RNA viruses could reflect infection patterns. The cases of synchrony likely imply that the community compositions changed in response to a varying third factor (e.g. through competition) or that dispersal introduced new organisms that caused community shifts^12^. In most cases, synchronous pairs were not significantly associated with a common third measure (Extended Data Fig. 3). The synchronous relationships most commonly observed here agree with a single-site marine study^23^; however, the diversity of sites presented here provide important counter examples to this emerging trend.

**Figure 2.**
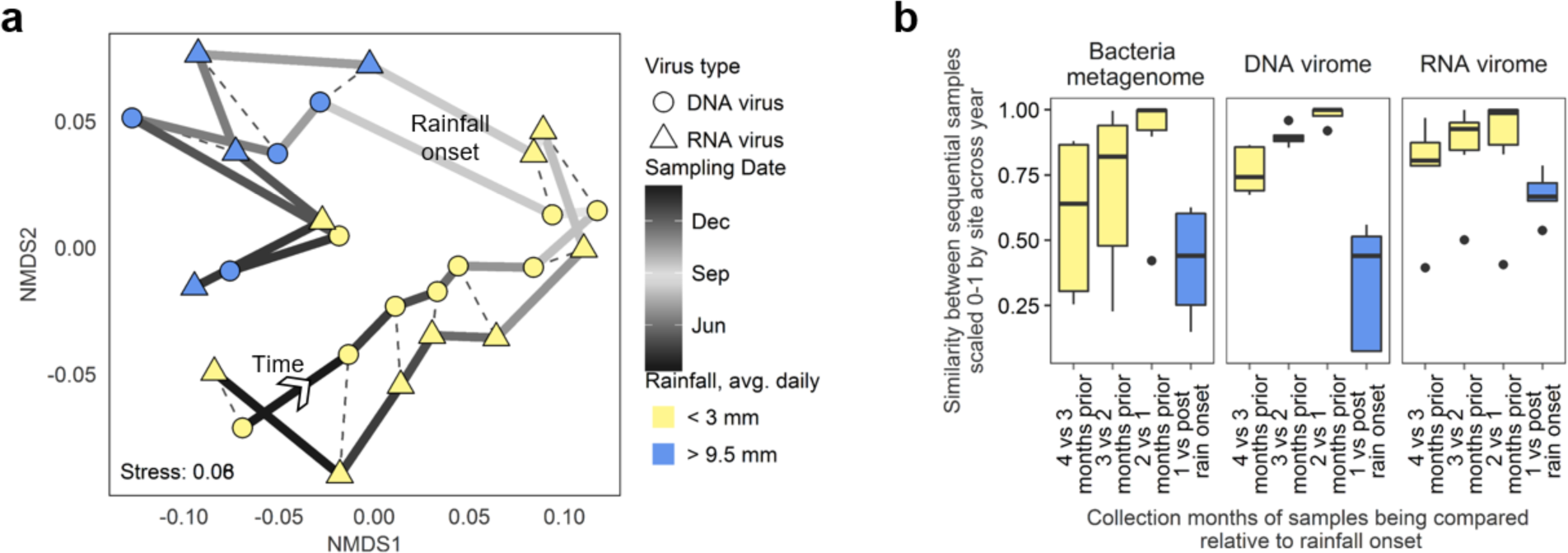
Onset of rainfall has consisent and large effect on riverine microplankton. **a**, NMDS plot of DNA & RNA viral communities from an agriculturally affected site (APL). Each point represents a viral community, solid lines connect sequential samples and are coloured by sampling date, dashed lines connect viromes extracted from the same sample. Points are coloured by the average rainfall over the three days prior to sampling. N= 13. **b**, Box plot of similarity between microbial communities collected in subsequent months, coloured by whether both sampling dates had low rainfall (yellow) or whether the earlier date was dry but later date had elevated rainfall (blue). N = 6 for bacteria and RNA viruses, N= 5 for DNA viruses.

Unexpectedly, DNA and RNA viral community compositions were synchronous in some sites (metagenomic and phylogenetic marker gene data, Mantel’s *r* = 0.4 - 0.6, q = 0.02 - 0.001), even though they were not consistently synchronous with bacteria or microeukaryotes (Extended Data Fig. 2). Because few, if any, studies have profiled DNA and RNA viral community compositions concurrently over time, this synchrony has not been previously investigated. While correlational data cannot prove the drivers of synchrony, environmental data can provide context. Synchronous DNA and RNA viromes were correlated with daylight hours (Extended Data Fig. 3) and a temporal trend is clear: sequential samples tended to be most alike and shift stepwise over time (Figure 2a, at one site; for other sites see Extended Data Fig. 4). This suggests that the DNA and RNA viral synchrony is not artefactual, but due to some temporal relationship, possibly with a common host group or synchronous groups.

Large shifts in DNA and RNA viromes in agriculturally affected sites were concurrent with the onset of rainfall after a dry period (Figure 2a, Extended Data Fig. 4). This trend was also observed in the other sampling sites and in bacterial communities (Figure 2b, microeukaryotic communities not tested due to insufficient data). These observations demonstrate the first-flush phenomenon; dry periods permit a buildup of solids, chemicals, metals, and organisms and the first significant rainfall causes an abrupt shift in the bacterial and viral communities in the receiving waters^24-26^. This shows that while continuous relationships with rainfall were not universal (Figure 1), response to a rainfall event was more common.

**Figure 3.**
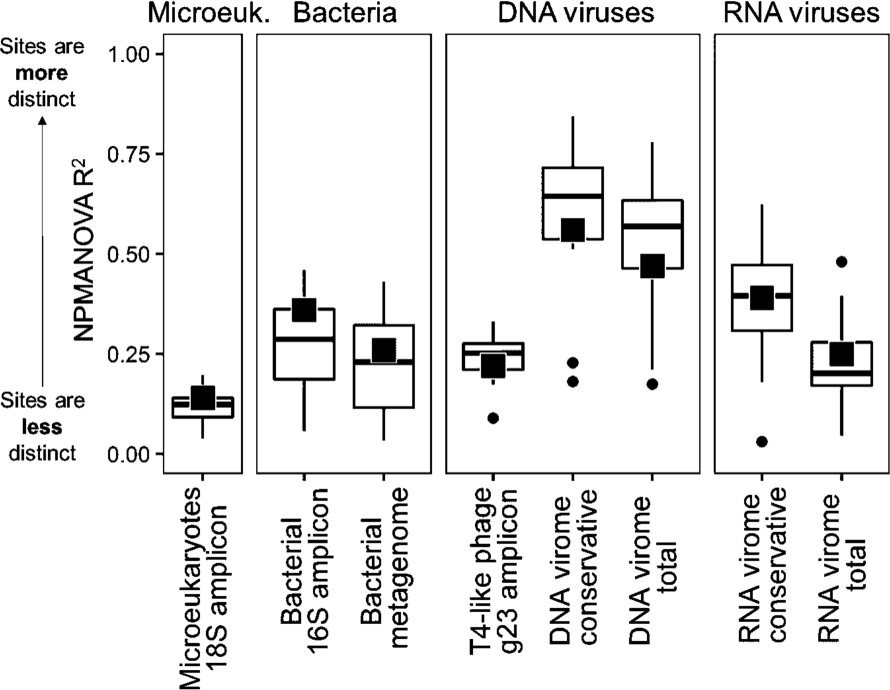
Geographic distinctiveness within viral, bacterial, and eukaryotic communities over 1 year of monthly samples. Proportion of variability among samples that is explained by sampling site (NPMANOVA R^2^), either across all sites (black square) or pairwise between sites (boxplots). In boxplots, the lower and upper box edges correspond to the first and third quartiles, the whiskers extend to the highest and lowest values that are within 1.5 times the inter-quartile range, and data beyond this limit are plotted as points.

While sampling site was a significant source of variation for all microbial groups, DNA viromes showed stronger geographic-based similarity than bacteria and microeukaryotes (Figure 3, Extended Data Fig. 1). This is consistent with the distinctiveness of T4-like bacteriophage seen in a study of polar lakes^27^. It is in contrast with the similarity of DNA viruses seen in two temperate lakes^28^, however these lakes are connected and have similar surrounding land use. Analysing bacterial amplicon data at a finer taxonomic resolution (99% identity OTUs) did not significantly increase its geographic distinctiveness (data not shown). This lower geographic distinctiveness of bacteria, particularly among sites with similar land use (pairwise NPMANOVA between the two agriculturally affected sites and between the two urban-affected sites: R^2^ <= 0.23, q = 0.0003, Extended Data Fig. 5), is consistent with previously shown low spatial stratification of bacteria among rivers^10-12^. In the one case where land use varied within a watershed (Figure 1a, AUP versus APL & ADS), land use and associated water chemistry differences appeared to override geographic proximity as a predictor of microbial community similarity (Extended Data Fig. 5). These findings support a major ecological role of dispersal at this geographic scale (10 – 130 km) for riverine bacterial and microeukaryotic plankton but reveals that viruses have a more distinct geographic pattern.

The higher geographic specificity of viruses observed here could reflect higher geographic specificity of host cells not sampled in this study, such as particle-associated plankton, riverbed biofilms, plants, humans, or other animals. Alternatively, viruses may be more geographically distinct because they replicate in the subset of microbial cells in the community that are active (estimated at 20-50% of bacterial cells^29^). This subset is more likely to be geographically distinct due to their increased susceptibility to selective pressures^29^ and more likely to be represented by viruses due to the mechanics of the lytic cycle and host-specificity^30^. Thus, we hypothesise that viruses may produce a stronger geographic signal than bacteria by amplifying the effect of species sorting against the background of widely dispersed inactive cells.

In conclusion, temporal and spatial profiling revealed contrasting patterns among superkingdoms and environmental conditions in riverine microbial plankton. Some relationships were common, such as microbial composition with day light hours and rainfall, and expected correlations were observed, such as between bacterial communities and pH. However, by examining multiple locations, these relationships were revealed not to be universal, even within similar sampling sites. This demonstrates the heterogeneity of riverine microbial ecosystems and the need for multi-site studies in riverine microbial ecology, as a similar study of a single site may have falsely concluded general trends. By examining multiple superkingdoms, correlations with nutrient concentrations were identified that would have been missed if only bacteria were profiled and the strong dispersal observed in bacteria and microeukaryotes was revealed not to extend to viruses. In summary, this study provides insight into the variability of microbiomes over superkingdoms, time, and space in an important, yet understudied environment. It reveals notable differences in community dynamics across microbial groups, and demonstrates the value of collectively studying microeukaryotes, bacteria and viruses across multiple time points and locations in microbiome studies.

## Methods

### Sampling & sequencing

River water was collected monthly for 12 to 13 consecutive months from six sites in three watersheds in southwestern British Columbia, Canada. The agricultural watershed had three sampling sites, one upstream of human activity (AUP), one adjacent to intensive agriculture (APL), and one further downstream (ADS). The urban watershed had two sampling sites, one with a catchment mix of forest and residential land use (UPL), and one further downstream with mostly residential and some park land use (UDS). The pristine watershed was in a protected forest area, with no land use (PUP). Sampling sites were not downstream of any lakes or dams. Water temperatures ranged from 3°C to 25°C. In the agricultural watershed, a distinct rainy period occurred from November to March, which is typical for the area. The other watersheds had more variable rainfall throughout the year. Sites from the same watershed were sampled on the same day. For full sampling and sequencing procedures see ^21^ and ^20^; a brief overview follows.

At each sampling event, 40 L of water was collected and then filtered sequentially to concentrate particles approximating the sizes of microeukaryotes (105 to 1 μm), bacteria (1 to 0.2 μm), and viral-sized particles^21^. Physical and chemical water measurements were also taken^20^. DNA was extracted from each size fraction, along with RNA from the viral-sized fraction^21^.

Amplicons for T4-like bacteriophages were prepared using primers targeting the myovirus g23 gene^21,31^. Amplicons for bacteria were prepared using primers targeting the V3-V4 regions of 16S rRNA gene^32,33^. Amplicons for microeukaryotes were prepared using primers targeting the V1-V3 regions of the 18S rRNA gene^34,35^. Amplicons were purified with a QIAQuick PCR Purification Kit (Qiagen Sciences, Maryland, MD) according to the manufacturer’s instructions. Sequencing libraries were prepared for amplicons using NEXTflex ChIP-Seq Kit (BIOO Scientific, Austin, TX), gel size-selected as per manufacturer’s instructions, and sequenced with 250-bp paired-end reads on an Illumina MiSeq platform (Illumina, Inc., San Diego, CA).

Bacterial metagenome libraries were prepared using Nextera XT DNA sample preparation kit (Illumina, Inc., San Diego, CA) and size selected using high-throughput gel-based Ranger technology^36^. Bacterial metagenomes were sequenced over multiple runs with 250 bp paired-end reads on an Illumina MiSeq, with positive controls (mock communities)^20,37^ and negative controls included in each run.

A modified adapter nonamer approach was used to synthesize viral cDNA and increase yields from the viral fraction^21,38^. Viral metagenome libraries were prepared from randomly amplified DNA and cDNA using NEXTflex ChIP-Seq kit (BIOO Scientific, Austin, TX) by following a gel-free option provided in the manufacturer’s instructions. These libraries were sequenced with 150 bp paired-end reads on an Illumina HiSeq platform (Illumina, Inc., San Diego, CA).

### DNA sequence pre-processing and quality control

Low quality bases were trimmed from the 3’ end of reads using a sliding window with a minimum Phred score of 20 (or 15 for g23) using Trimmomatic^39^. Adapters were removed using Cutadapt^40^ with default parameters. Paired-end reads were merged using PEAR^41^. Microeukaryotic 18S amplicon paired-end reads could not be merged, so Operational Taxonomic Units (OTUs) were generated from reads with the same primer sequence.

T4-like myovirus g23 amplicons reads were translated into amino acid sequences using Fraggenescan v1.16 with the Illumina 5% error model (Rho, Tang, and Ye 2010). OTUs were generated using USEARCH^42^ v7: sequences were dereplicated, clustered at 95% identity, then all reads were mapped back against cluster representatives to calculate abundances. Sample read totals were subsampled to 10,000 reads using the vegan package^43^ in R^44^ v3.1.2. Random resampling was performed 10,000 times and the median value of all iterations was chosen. Bacterial 16S and microeukaryotic 18S OTUs were generated from amplicon reads using the Mothur^45^ MiSeq clustering protocol^46^ and rarefied to 10,000 reads.

Metagenomic reads were trimmed at the 3’ end with a sliding window with a minimum Phred score of 20 using Trimmomatic^39^. DNA virome reads shorter than 70 bp were discarded, resulting in a dataset of 20 Gb in 225 M reads. RNA virome reads shorter than 100 bp were discarded and ribosomal reads were removed using meta-rRNA^47^, resulting in a dataset of 17 Gb across 149 M reads. Bacterial metagenome reads shorter than 100 bp were discarded, resulting in a dataset of 16 Gbp across 75 M reads.

### Generation of high-confidence DNA & RNA virome datasets

Viromes were assembled using CLC and proteins were predicted from contigs using Prodigal in metagenomic mode with default parameters. Predicted proteins at least 26 amino acids long were clustered *de novo* using parallel cd-hit^48^, with criteria as previously used^49^: word length of 4 and 60% identity over 80% length of the shorter sequence. Reads were assigned to clusters with a blastx-style similarity search against cluster representative sequences using DIAMOND^50^ with minimum 60% sequence similarity over minimum 26 amino acid alignment length. While protein cluster analysis is common in large scale marine studies^49,51^, we did not use this dataset for primary analysis as many samples had a small proportion of reads in any protein cluster (mean 13%, range 8-30% of DNA virus reads and mean 25%, range 8-60% for RNA virus reads).

Contigs were tested for amino acid sequence similarity to reference sequences in NCBI’s nr database using RAPSearch and taxonomically classified using MEGAN5. A small proportion of contigs were assigned as DNA viral (4% of contigs, 0.7% of total reads) and RNA viral (2% of contigs, 7% of total reads).

In the DNA virome dataset, 42% of contigs were assigned as bacterial, corresponding to 20% of assembled reads and 7% of total reads. To assess whether these bacterial assignments were due to miss-assignment of viral sequences (e.g. auxiliary metabolic genes, prophages) or an indication of bacterial contamination (e.g. from laboratory reagents, free-floating DNA, or host DNA packaged in viral capsid)^52^, reads were tested for the presence of bacterial genes unlikely to occur in viruses. Across 515,000-read subsets of samples, similarity to the 16S rRNA gene was found in 1 to 156 reads (mean: 30, standard deviation: 25). Though these are small numbers, they are an indication of the number of bacterial genomes potentially present. This means that the contigs identified as bacterial in the taxonomic results cannot be ruled out as bacterial contamination. Further, the contigs that were left unassigned by the taxonomic classification also cannot be ruled out as bacterial.

To remove potential bacterial contamination from the DNA and RNA viromes, subsets of the read data were generated that only included sequences from protein clusters with at least one member that was assigned as coming from DNA or RNA viruses, respectively. This reduced the number of reads per sample from 515,000 in the “total” dataset to 10,000 in the “conservative” subset for DNA viromes and from 45,000 to 1,000 for RNA viromes. As this is a fairly small number of reads, we estimated the stability of distance matrices with low numbers of reads (see below) and used both total and conservative datasets to test trends.

### Sample similarity estimation & spatiotemporal analysis

Pairwise similarity between amplicon samples was performed using vegan^43^ in R^44^ to calculate Bray-Curtis dissimilarity between OTU abundance profiles. Pairwise similarity between metagenomes was assessed using Mash distances v1.0.2^22^, which compares metagenomes based on k-mer presence-absence. For display in heatmaps in Extended Data Fig. 1, extreme values of similarities were collapsed to be represented by one color. Extreme values were defined as those values more than 2.5 times the median absolute deviation (MAD) away from the median^53^. Collapsed values were only used for display and not for any statistical tests.

Due to the small number of reads in the conservative RNA virus dataset, we investigated whether this depth was enough to obtain a stable representation of the communities. We randomly selected 1,000 reads ten times per sample from 68 samples which had at least 10,000 reads in the conservative RNA virus dataset. We ran Mash on these subsamples and calculated the pairwise Mantel correlations between the resultant dissimilarity matrices. All matrices had correlation scores of at least R = 0.95 with Pearson’s correlation and R=0.94 with Spearman’s correlation. We decided this consistency was sufficiently high to justify confidence in high level patterns within this data.

All statistical tests were performed in R^44^ v3. Permutation-based p values were calculated using 9999 permutations. Multiple test correction was performed where appropriate using the Benjamini-Hochberg procedure and adjusted p values reported as q values. Significance test values were considered statistically significant if lower than 0.05, except where indicated otherwise.

The proportion of variability among sample similarities that could be explained by sampling site was estimated using NPMANOVA as implemented in the adonis function from the vegan R package^43^. Gene family variability was based on SEED subsystem classifications^20^ and calculated using Bray-Curtis dissimilarities. The NMDS plots in Figure 2 and Extended Data Fig. 4 were generated using the vegan metaMDS function, with rotation and scaling of ordinations performed using the procrustes function and tested for significance using the pro.test function. Samples from April 2013 (105 and 106) were highly dissimilar and removed from Figure 2a to permit the trend in the other 12 samples to be displayed.

Synchrony was tested using Mantel matrix correlation tests with Spearman correlations, implemented in the vegan R package^43^. When testing samples from multiple sites for synchrony, a partial Mantel test was used to control for geographic distance between sampling sites. Environmental data were tested for correlations with microbial community similarities using the envfit function. If applicable, the environmental measures to test were selected based on their magnitude and variability in the context of water quality guidelines^54^. Relationships among environmental measures were assessed using Spearman’s correlation. Correlations within and among environmental measures and microbial community similarities were displayed in a network using the visNetwork R package. Correlations that had a q value less than 0.1 were considered statistically significant. Correlations that had a q value greater than 0.1 but a p value less than 0.05 were not considered statistically significant but were included in visualisations to avoid overconfidence in the absence of a relationship, however, they should be interpreted with caution.

## Data Availability

All raw sequences are deposited in the NCBI Sequence Read Archive under BioProject accession PRJNA287840.

## Acknowledgements

We thank Jared R. Slobodan and Matthew J. Nesbitt from Coastal Genomics Inc., Burnaby, BC, Canada for their assistance in applying Ranger Technology for DNA sequencing library size selection. This work was funded by Genome BC and Genome Canada grant No. LSARP-165WAT, with major support from the Simon Fraser University Community Trust Endowment Fund and additional support from the Public Health Agency of Canada.

## Author contributions

J.I.R, P.K.C.T., N.A.P., C.A.S, and F.S.L.B. designed the study, guided the analyses, aided in interpretations, and acquired funding. M.I.U. led the sampling and sequencing, with assistance from M.C., M.A.C, and K.I.C̤ J.R.S. and M.J.N. performed size selection of sequencing libraries. T.V. led the bioinformatics and data analysis and wrote the manuscript, with significant input from F.S.L.B. A.T. compiled OTU tables for the g23 data. M.V., M.A.P., and W.W.L.H. guided analyses. All authors contributed to final revisions of the manuscript.

## References

1. Baird, D. J. & Hajibabaei, M. Biomonitoring 2.0: A new paradigm in ecosystem assessment made possible by next-generation DNA sequencing. Mol. Ecol. 21, 2039–2044 (2012).

2. Meybeck, M. Global analysis of river systems: from Earth system controls to Anthropocene syndromes. Philos. Trans. R. Soc. Lond. B. Biol. Sci. 358, 1935–55 (2003).

3. Findlay, S. Stream microbial ecology. J. North Am. Benthol. Soc. 29, 170–181 (2010).

4. Zeglin, L. H. Stream microbial diversity in response to environmental changes: review and synthesis of existing research. Front. Microbiol. 6, 454 (2015).

5. Middelboe, M., Jacquet, S. & Weinbauer, M. Viruses in freshwater ecosystems: An introduction to the exploration of viruses in new aquatic habitats. Freshw. Biol. 53, 1069–1075 (2008).

6. Peduzzi, P. Virus ecology of fluvial systems: a blank spot on the map? Biol. Rev. 91, 937–949 (2016).

7. Jacquet, S., Miki, T., Noble, R., Peduzzi, P. & Wilhelm, S. Viruses in aquatic ecosystems: important advancements of the last 20 years and prospects for the future in the field of microbial oceanography and limnology. Adv. Oceanogr. Limnol. 1, 97–141 (2010).

8. Ruiz-González, C., Niño-García, J. P., Lapierre, J.-F. & del Giorgio, P. A. The quality of organic matter shapes the functional biogeography of bacterioplankton across boreal freshwater ecosystems. Glob. Ecol. Biogeogr. n/a-n/a (2015). doi:10.1111/geb.12356

9. Staley, C. et al Species sorting and seasonal dynamics primarily shape bacterial communities in the Upper Mississippi River. Sci. Total Environ. 505, 435–445 (2015).

10. Niño-García, J. P., Ruiz-González, C. & del Giorgio, P. A. Interactions between hydrology and water chemistry shape bacterioplankton biogeography across boreal freshwater networks. ISME J. 1–12 (2016). doi:10.1038/ismej.2015.226

11. Jackson, C. R., Millar, J. J., Payne, J. T. & Ochs, C. A. Free-living and particle-associated bacterioplankton in large rivers of the Mississippi River basin demonstrate biogeographic patterns. Appl. Environ. Microbiol. 80, 7186–7195 (2014).

12. Crump, B. C. et al Circumpolar synchrony in big river bacterioplankton. Proc. Natl. Acad. Sci. U. S. A. 106, 21208–12 (2009).

13. Read, D. S. et al Catchment-scale biogeography of riverine bacterioplankton. ISME J. 9, 516–526 (2014).

14. Thomas, M. C., Selinger, L. B. & Inglis, G. D. Seasonal diversity of planktonic protists in Southwestern Alberta rivers over a 1-year period as revealed by terminal restriction fragment length polymorphism and 18S rRNA gene library analyses. Appl. Environ. Microbiol. 78, 5653–5660 (2012).

15. Bradford, T. M. et al Microeukaryote community composition assessed by pyrosequencing is associated with light availability and phytoplankton primary production along a lowland river. Freshw. Biol. 58, 2401–2413 (2013).

16. Simon, M. et al Marked seasonality and high spatial variability of protist communities in shallow freshwater systems. ISME J. (2015). doi:10.1038/ismej.2015.6

17. Dann, L. M. et al Marine and giant viruses as indicators of a marine microbial community in a riverine system. Microbiologyopen (2016). doi:10.1002/mbo3.392

18. Silva, B. S. et al Virioplankton Assemblage Structure in the Lower River and Ocean Continuum of the Amazon. mSphere 2, e00366–17 (2017).

19. Aguirre de Cárcer, D., López-Bueno, A., Pearce, D. A. & Alcamí, A. Biodiversity and distribution of polar freshwater DNA viruses. Sci. Adv. 1, e1400127 (2015).

20. Van Rossum, T. et al Year-Long Metagenomic Study of River Microbiomes Across Land Use and Water Quality. Front. Microbiol. 6, 1405 (2015).

21. Uyaguari-Diaz, M. I. et al A comprehensive method for amplicon-based and metagenomic characterization of viruses, bacteria, and eukaryotes in freshwater samples. Microbiome 4, 20 (2016).

22. Ondov, B. D. et al Mash: Fast genome and metagenome distance estimation using MinHash. Genome Biol. 29827 (2015). doi:10.1101/029827

23. Chow, C.-E. T., Kim, D. Y., Sachdeva, R., Caron, D. A. & Fuhrman, J. A. Top-down controls on bacterial community structure: microbial network analysis of bacteria, T4-like viruses and protists. ISME J. 8, 816–29 (2014).

24. Williamson, K. E., Harris, J. V., Green, J. C., Rahman, F. & Chambers, R. M. Stormwater runoff drives viral community composition changes in inland freshwaters. Front. Microbiol. 5, (2014).

25. Deletic, A. The first flush load of urban surface runoff. Water Res. 32, 2462–2470 (1998).

26. Tseng, C.-H. et al Microbial and viral metagenomes of a subtropical freshwater reservoir subject to climatic disturbances. ISME J. 7, 2374–2386 (2013).

27. De Cárcer, D. A., Pedrós-Alió, C., Pearce, D. A. & Alcamí, A. Composition and interactions among bacterial, microeukaryotic, and T4-like viral assemblages in lakes from both polar zones. Front. Microbiol. 7, (2016).

28. Mohiuddin, M. & Schellhorn, H. Spatial and temporal dynamics of virus occurrence in two freshwater lakes captured through metagenomic analysis. Front. Microbiol. 6, (2015).

29. Lennon, J. T. & Jones, S. E. Microbial seed banks: the ecological and evolutionary implications of dormancy. Nat. Rev. Microbiol. 9, 119–130 (2011).

30. Paez-Espino, D. et al Uncovering Earth’s virome. Nature 536, 425–430 (2016).

31. Filée, J., Tétart, F., Suttle, C. A. & Krisch, H. M. Marine T4-type bacteriophages, a ubiquitous component of the dark matter of the biosphere. Proc. Natl. Acad. Sci. U. S. A. 102, 12471–6 (2005).

32. Muyzer, G., de Waal, E. C. & Uitterlinden, A. G. Profiling of complex microbial populations by denaturing gradient gel electrophoresis analysis of polymerase chain reaction-amplified genes coding for 16S rRNA. Appl. Environ. Microbiol. 59, 695–700 (1993).

33. Caporaso, J. G. et al Global patterns of 16S rRNA diversity at a depth of millions of sequences per sample. Proc. Natl. Acad. Sci. U. S. A. 108 Suppl, 4516–22 (2011).

34. Zhu, F., Massana, R., Not, F., Marie, D. & Vaulot, D. Mapping of picoeucaryotes in marine ecosystems with quantitative PCR of the 18S rRNA gene. FEMSMicrobiol. Ecol. 52, 79–92 (2005).

35. Amann, R. I., Krumholz, L. & Stahl, D. A. Fluorescent-oligonucleotide probing of whole cells for determinative, phylogenetic, and environmental studies in microbiology. J. Bacteriol. 172, 762–770 (1990).

36. Uyaguari-Diaz, M. I. et al Automated Gel Size Selection to Improve the Quality of Next-generation Sequencing Libraries Prepared from Environmental Water Samples. J. Vis. Exp. e52685 (2015). doi:10.3791/52685

37. Peabody, M. A., Van Rossum, T., Lo, R. & Brinkman, F. S. L. Evaluation of shotgun metagenomics sequence classification methods using in silico and in vitro simulated communities. BMC Bioinformatics 16, 363 (2015).

38. Wang, D. et al Microarray-based detection and genotyping of viral pathogens. Proc. Natl. Acad. Sci. U. S. A. 99, 15687–92 (2002).

39. Bolger, A. M., Lohse, M. & Usadel, B. Trimmomatic: a flexible trimmer for Illumina sequence data. Bioinformatics 30, 2114–20 (2014).

40. Martin, M. Cutadapt removes adapter sequences from high-throughput sequencing reads. EMBnet.journal 17, 10–12 (2011).

41. Zhang, J., Kobert, K., Flouri, T. & Stamatakis, A. PEAR: a fast and accurate Illumina Paired-End reAd mergeR. Bioinformatics 30, 614–20 (2014).

42. Edgar, R. C. Search and clustering orders of magnitude faster than BLAST. Bioinformatics 26, 2460–1 (2010).

43. Oksanen, J. et al Package ‘vegan’: Community Ecology Package. Community ecology package, version 2, 280 (2015).

44. R Core Team. R: A language and environment for statistical computing. (2013).

45. Schloss, P. D. et al Introducing mothur: open-source, platform-independent, community-supported software for describing and comparing microbial communities. Appl. Environ. Microbiol. 75, 7537–41 (2009).

46. Kozich, J. J., Westcott, S. L., Baxter, N. T., Highlander, S. K. & Schloss, P. D. Development of a dual-index sequencing strategy and curation pipeline for analyzing amplicon sequence data on the miseq illumina sequencing platform. Appl. Environ. Microbiol. 79, 5112–5120 (2013).

47. Huang, Y., Gilna, P. & Li, W. Identification of ribosomal RNA genes in metagenomic fragments. Bioinformatics 25, 1338–1340 (2009).

48. Fu, L., Niu, B., Zhu, Z., Wu, S. & Li, W. CD-HIT: accelerated for clustering the next-generation sequencing data. Bioinformatics 28, 3150–2 (2012).

49. Hurwitz, B. L. & Sullivan, M. B. The Pacific Ocean Virome (POV): A Marine Viral Metagenomic Dataset and Associated Protein Clusters for Quantitative Viral Ecology. PLoS One 8, e57355 (2013).

50. Buchfink, B., Xie, C. & Huson, D. H. Fast and sensitive protein alignment using DIAMOND. Nat. Methods 12, 59–60 (2014).

51. Brum, J. R. et al Patterns and Ecological Drivers of Ocean Viral Communities. Science (80-.). 348, 1261498-1–11 (2015).

52. Hurwitz, B. L., U’Ren, J. M. & Youens-Clark, K. Computational prospecting the great viral unknown. FEMS Microbiology Letters 363, (2016).

53. Leys, C., Ley, C., Klein, O., Bernard, P. & Licata, L. Detecting outliers: Do not use standard deviation around the mean, use absolute deviation around the median. J. Exp. Soc. Psychol. 49, 764–766 (2013).

54. Canadian Council of Ministers of the Environment. Canadian Environmental Quality Guidelines and Summary Table. (2007).

